# Single-cell RNA sequencing in Hirschsprung’s disease tissues reveals lack of neuronal differentiation in the aganglionic colon segment

**DOI:** 10.1101/2025.07.01.662516

**Authors:** Szabolcs Tarapcsak, Xiaomeng Huang, Yi Qiao, Andrew Farrell, Lija Mammen, Amy Lovichik, Gayatri D. Khanderao, Teresa Musci, Philip J Moos, Matthew A Firpo, Michael Rollins, Gabor T Marth

**Affiliations:** Eccles Institute of Human Genetics, University of Utah, Salt Lake City, Utah, USA; Division of Pediatric Surgery, University of Utah Health, Salt Lake City, Utah, USA; Department of Pathology, Primary Children’s Hospital, Salt Lake City, Utah, USA; Department of Surgery, University of Utah School of Medicine, Salt Lake City, Utah, USA; Department of Pharmacology and Toxicology, College of Pharmacy, University of Utah, Salt Lake City, Utah, USA

## Abstract

The enteric nervous system (ENS) is a complex network of neurons and glial cells. Hirschsprung’s disease (HSCR) is a congenital condition characterized by the absence of ganglion cells in the distal colon, leading to functional bowel obstruction. In this study, we used single-cell RNA sequencing (scRNA-seq) and whole genome sequencing (WGS) to analyze healthy and aganglionic colon segments from HSCR patients. Using scRNA-seq, we identified 13 major cell types in patient samples and observed that neural progenitor cells were present in both healthy and aganglionic colon regions, while mature neurons were absent from aganglionic colon. In these progenitor cells, critical differentiation pathway genes displayed reduced expression in the aganglionic colon, suggesting a disruption in their transition to mature neuronal cell types. Furthermore, transcriptomic analysis revealed significant alterations in gene expression across several stromal cell types. These transcriptomic shifts, particularly in mast cells, support the hypothesis that altered gene expression in the microenvironment of neural progenitor cells contributes to impaired differentiation. Our findings support the hypothesis that neural precursors in HSCR are capable of migration, but they are defective in their differentiation to mature cell types. Our analysis provides insights into potential therapeutic targets to stimulate neurogenesis in the aganglionic colon.

## Introduction

The enteric nerve system (ENS) is a complex network of mature neurons and glial cells. The ENS is the intrinsic nervous system of the gastrointestinal tract that is responsible for the regulation of sophisticated biological processes, including digestion and absorption, secretion of gastrointestinal hormones and signaling molecules, preservation of the immune cell repertoire and maintenance of the healthy microbiome. ENS is comprised of app. 200-600 million neurons and seven times more glial cells (1) that control these biological processes without signaling input from the central nervous system (2).

Hirschsprung’s disease (HSCR) is the best known congenital disorder of the enteric nervous system (ENS), affecting 1 in 5000 individuals. The hallmark of HSCR is the lack of ganglion cells in a variable length of the distal part of the colon (aganglionosis) (3). The concomitant absence of mature enteric neurons in the affected colon segment leads to a tonic contraction along with no propulsive motility. These lead to gastrointestinal obstruction, chronic constipation, abdominal distention, vomiting and growth failure (3, 4). Treatment for HSCR is based on the removal of the aganglionic colon section, but in many cases HSCR patients continue to have gastrointestinal symptoms even after surgery (5, 6). Despite advancements in surgical treatments, many patients experience persistent bowel dysfunction, highlighting the need for further understanding of HSCR pathogenesis at the cellular and molecular levels.

HSCR is understood to be a defect in developmental processes. ENS development is a complex biological process that is controlled by many regulatory mechanisms. The ENS is derived from the neural crest (NC), an embryonic tissue populated by highly proliferative cell populations showing high mobility (7). NC-derived cells (enteric neural crest derived cells, ENCCs) migrate through the mesenchyme and populate the length of the bowel in a protracted process (between embryonic week 3 and 7 in humans) (7, 8). ENS is populated primarily by vagal-derived ENCCs and to a smaller extent by sacral-derived ENCCs (9). Recent studies also showed that other cells, including Schwann cell precursors (SCPs) provide further precursor pools for the formation of the ENS. These cells primarily reach the gut through extrinsic nerve fibers (10). Parallel with their migration, ENCCs proliferate extensively and in humans they undergo inward radial migration, forming the two layers of neural cells that comprise the myenteric and submucosal plexuses (11). Subsequently ENCCs differentiate into mature neurons and glia and form a network of ganglia throughout the bowel (12). Hindered migration, proliferation and/or differentiation of ENCCs therefore can lead to HSCR. The exact mechanism regulating the journey of ENCCs is not well-understood, but many signaling pathways, e.g. the *RET/GFRα1/GDNF*, the *EDNRB/ECE1/EDN3*, sonic hedgehog and *NOTCH* pathways and transcription factors regulating the *RET/EDNRB* pathways, e.g. *SOX10* and *PHOX2B* are involved (13, 14)(15).

Only about 20% of HSCR cases show family history with a Mendelian type of inheritance while sporadic cases show a non-mendelian type of inheritance involving various genetic and environmental factors (16). Several genes have been associated with ENS precursor migration and differentiation contributing to the pathogenesis of HSCR. Most importantly, a transmembrane tyrosine kinase receptor *RET* has been identified as the main HSCR-associated gene: *RET* is mutated in 50% of familial and 20% of sporadic HSCR cases (17). Using linkage mapping and mouse models several further HSCR susceptibility genes were identified that are involved in the development of the ENS including *EDNRB, GDNF, ECE1, GFRα1, NRG1, PHOX2B, NRTN, TCF4, SOX10, EDN3, NTN, PSPN, ZFHX1B, L1CAM* and *KIAA1279* (*18–20*). However, such genes associated with HSCR susceptibility can explain only 30% of total HSCR incidence (21) suggesting the involvement of other, low-penetrance gene mutations and transcriptional changes in ENCCs and stromal cells in HSCR pathogenesis. Interestingly, in HSCR mouse models (Ret-/-) it has been observed that the transcriptional profile of gut mesenchyme and epithelium differs from that of wild-type mice (22). Together with the discovery that identical genetic backgrounds can produce aganglionic colons of different lengths, these findings highlight the crucial role of transcriptomic alterations in the tissue environment as a driving factor in HSCR etiology (23).

Single cell RNA sequencing (scRNA-seq) is a powerful technique to compare the transcriptomic profile of different cell-types and for the identification of rare cellular populations or intermediate states in tissue samples. Of note, scRNA-seq has been used in many studies portraying the importance of transcriptional changes in gut development and pathobiology, as summarized in (24). There is only a limited number of studies using scRNA-seq for analyzing the ENS (25). It has been shown that in embryonic gut neural crest cells differentiate in two early types of neurons marked by *Etv1* and *Bnc2* expression (26) and later these progenitor cells differentiate into 11 distinct neural subtypes (27). Moreover, scRNA-seq has been utilized in studies aiming to characterize the transcriptomic alterations specifically in HSCR. In a recent study, expression of HSCR-associated genes were analyzed in human embryonic gut tissue samples using scRNA-seq. In this study, key differences in the expression profiles between differentiating cell populations and gut regions were identified (26). More recently scRNAseq has been used to analyze early ENS progenitor cells differentiated from pluripotent stem cells to identify vagal NC subpopulations (28).

Despite these advances, a comprehensive understanding of the transcriptomic changes occurring in both the healthy and aganglionic segments of the colon in HSCR patients is lacking. No study to date has systematically compared the transcriptomes of multiple cell types from both healthy and diseased tissues at a single-cell resolution. Our research aims to address this gap by performing combined scRNA-seq and whole genome sequencing (WGS) on the proximal (healthy) and distal (aganglionic) colon segments of HSCR patients.

## Results

### Single-cell transcriptomic landscape of healthy and aganglionic colon in HSCR patients

We sampled the aganglionic and healthy segments of the colon from three HSCR patients and collected cell suspensions from the tissue samples. Colon tissues were dissociated into single cell suspensions and scRNA-seq experiments were performed. Simultaneously, DNA samples were collected from tissue samples and matched blood for WGS analysis. Using collected WGS data, we have shown that several well-known HSCR-associated genes harbored germline variants in our patient cohort, including *RET, DHCR7, TCF4 and NRG3* (**Supplementary table 1**). Many of these are well-known pathogenic variants described in the ClinVar database, however 35% of identified missense variants in these genes were not included in ClinVar (**Supplementary table 1**).

Based on the collected scRNA-seq data of three patients, overall 20,490 cells (7,647 healthy- and 12,843 aganglionic sample cells) were analyzed. Following dimension reduction of the data using UMAP (**Fig. 1A, D**) we analyzed the transcriptomic profile of the cell clusters (**Fig. 1B, C**) and identified 13 major cell types in collected HSCR colon samples using specific marker gene sets (see **Supplementary table 2**). Cell type ratios of aganglionic and healthy colon samples showed no major differences (paired T-test) (**Fig. 1E**), with the exception of mature neurons, that were only present in the healthy colon samples (healthy colon: 0.17±0.01%, aganglionic colon: 0±0.01%, p<0.0001, T-test), however large variability was observed in the calculated ratios of these cells between patients (**Fig. 1E**) representing sampling variability rather than relevant biological differences.

**Figure 1:**
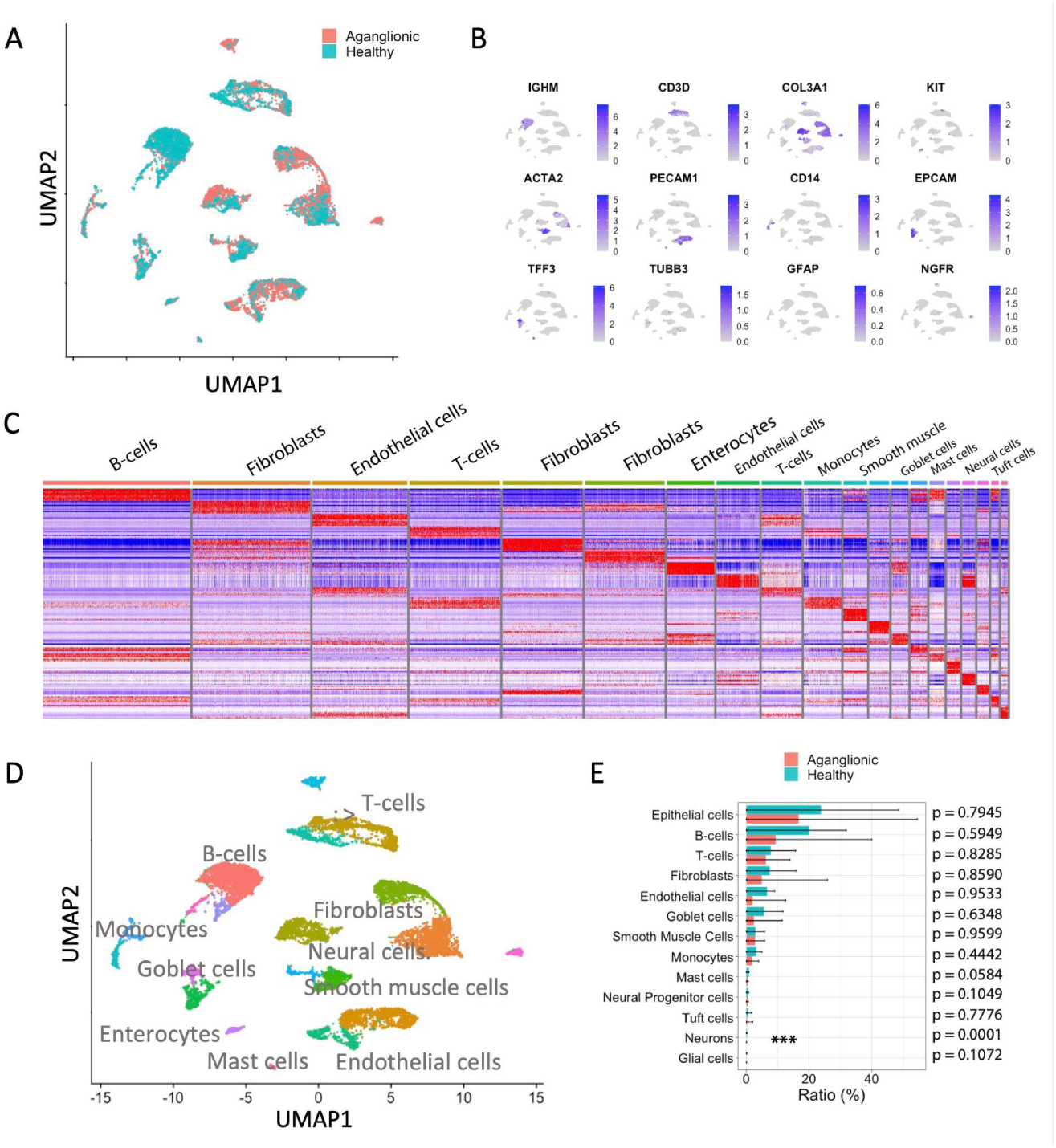
Experimental design and overview of scRNA-seq analysis of HSCR patient samples. (**A**) Representative UMAP embedding overlay showing the single cell transcriptome distribution of healthy and aganglionic colon samples. (**B**) Feature plots of representative marker gene expressions used for cell type identification. (**C**) Heatmap of top 10 most significantly upregulated genes of identified cell clusters in a representative HSCR patient. (**D**) Representative UMAP embedding of single cell transcriptomes from HSCR healthy and aganglionic colon samples with cell type annotations. (**E**) Calculated ratios of cell-types in the healthy and aganglionic colon samples (n=3, mean±SD, T-test).

Our scRNA-seq analysis suggested that, aside from the absence of mature neurons, Hirschsprung’s disease does not induce widespread changes in the overall cellular composition of the aganglionic colon segment relative to the healthy colon.

### Neural progenitor cells identified in the aganglionic colon of HSCR patients

Given that the primary hallmark of Hirschsprung’s disease (HSCR) is the absence of differentiated neurons in the aganglionic colon, we sought to comprehensively characterize the neural progenitor and neural cell populations in both healthy and aganglionic colon segments from HSCR patient samples.

Using specific marker genes (see **Supplementary table 2**), we identified mature neurons, glial cells, and neural progenitor cells by isolating a subset of cells exhibiting neural-like expression patterns (**Fig. 2A**). Our analysis confirmed the absence of mature neurons in the aganglionic colon segments (healthy colon: 0.17±0.01%, aganglionic colon: 0±0.01%, p<0.0001, T-test) (**Fig. 2B**). However, we detected a small population of neural progenitor cells (*NGFR+, UCHL1-, S100B-*) in both healthy and aganglionic colon segments, although at slightly (but not statistically significantly) lower numbers in the aganglionic region compared to the healthy colon (healthy colon: 0.65±0.05%, aganglionic colon: 0.45±0.16%, p=0.1049, T-test) (**Fig. 2B, C**). These progenitor cells lacked expression of mature neuron markers (*TUBB3, UCHL1, GAL*) and mature glial cell markers (*S100B, GFAP, ERBB3*), indicating that they remained in an undifferentiated state (**Fig. 2C**). To validate our scRNA-seq findings, we performed immunofluorescence microscopy, which confirmed the presence of p75 (neurotrophin receptor, marker of neural progenitor cells, encoded by *NGFR*) protein-expressing cells in both healthy and aganglionic colon segments (**Fig. 2D**).

**Figure 2:**
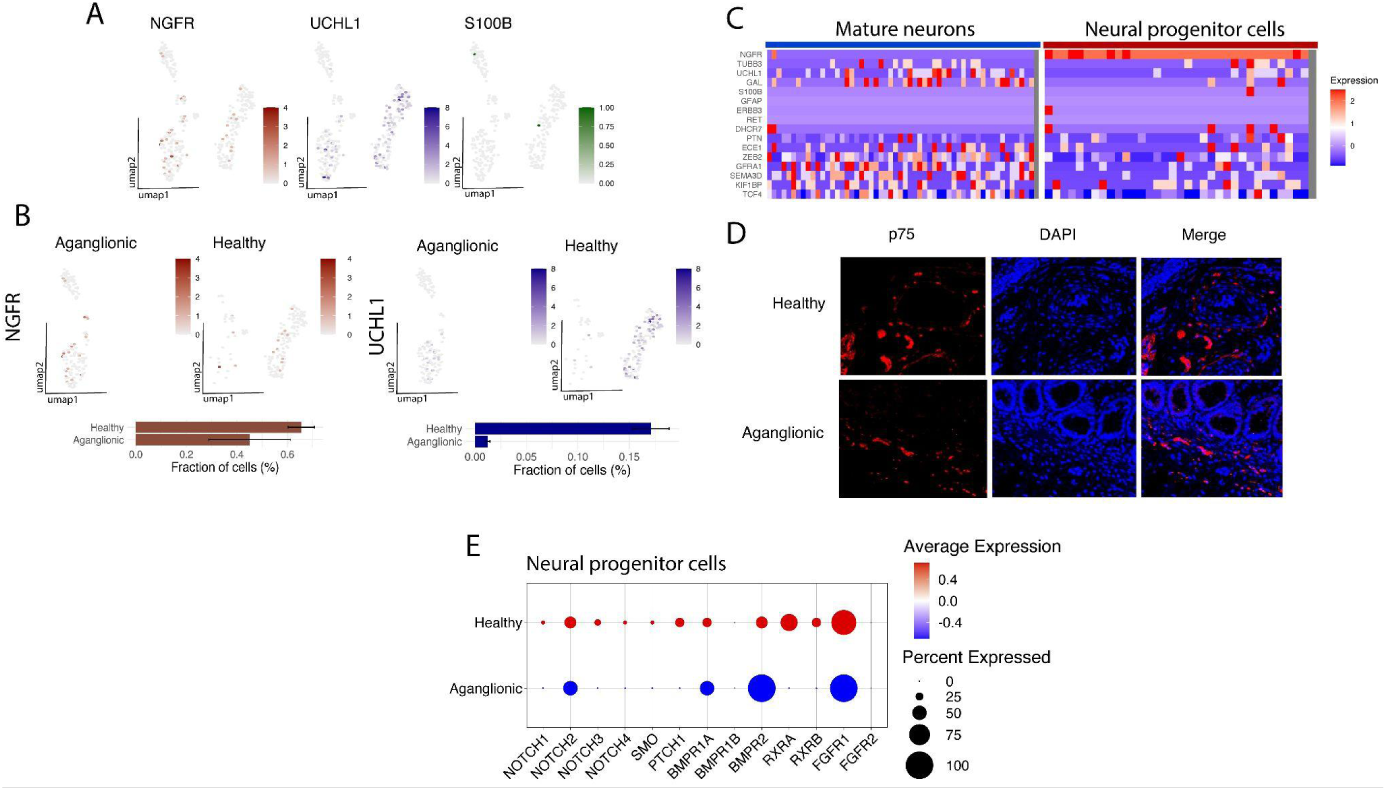
Identification of neural-progenitor cells in the aganglionic colon of HSCR patient. (**A**) UMAP plots showing expression of neural progenitor cell- (NGFR), mature neuron (UCHL1) and mature glial cell (S100B) marker gene expression in a representative patient sample. (**B**) UMAP plots showing expression of neural progenitor cell- (NGFR) and mature neuron (UCHL1) marker gene expression in a representative patient sample split by tissue origin. Barcharts show mean+SD of neural progenitor and mature neuron cell ratios of three HSCR patients. (**C**) Heatmap showing the expression of neural progenitor, mature neuron and mature glial cell markers and HSCR-associated genes in identified mature neurons and neural progenitor cells. (**D**) Fluorescent microscopic images of p75 (*NGFR*) and DAPI stained colon samples from HSCR patients show the presence of neural progenitor cells in the healthy and the aganglionic colon samples alike. (**E**) Dotplot of gene expression of key differentiation pathway receptor genes expressed on healthy and aganglionic colon segment resident neural progenitor cells (representative plot).

We next analyzed the expression of known HSCR-associated genes in both mature neurons and neural progenitor cells. Mature neurons displayed elevated expression of key HSCR-associated genes, including *GFRA1, ZEB2, TCF4* and *ECE1* (**Fig. 2C**). In contrast, these genes were expressed at significantly lower levels in neural progenitor cells (e.g., *GFRA1, SEMA3D, ECE1* and *TCF4*) (**Fig. 2C**). Notably, no *RET* expression was observed in either the neural progenitor cells or mature neurons from any of the three patient samples analyzed (**Fig. 2C**). Furthermore we compared the gene expression profile of healthy-aganglionic colon resident neural progenitor cells in our patient cohort and showed that for several differentiation factor receptor genes, significantly lower expression levels were observed in the neural progenitor cells of the aganglionic colon (e.g. *NOTCH1/3, SMO, PTCH1, RXRA, RXRB*) as compared to the healthy colon resident neural progenitor cells (**Fig. 2E**).

Our findings demonstrate that neural progenitor cells are present in the aganglionic colon segments of HSCR patients and critical genes in the *NOTCH* and *SHH* differentiation pathway displayed reduced expression in these cells supporting the hypothesis that neural precursors in HSCR are capable of migration, but they are defective in their differentiation to mature cell types.

### Potential link between transcriptomic changes in stroma cells and impaired neural differentiation in aganglionic colon cells in HSCR

Analysis of scRNA-seq data from healthy and aganglionic colon segments revealed differentially expressed genes across multiple cell types, including epithelial and immune cells (**Table 1**). This analysis focused on genes consistently altered across all HSCR patients. Our findings demonstrated significant downregulation of numerous genes in various cell populations within the aganglionic colon segments (**Fig. 3**, **Table 1**).

**Figure 3:**
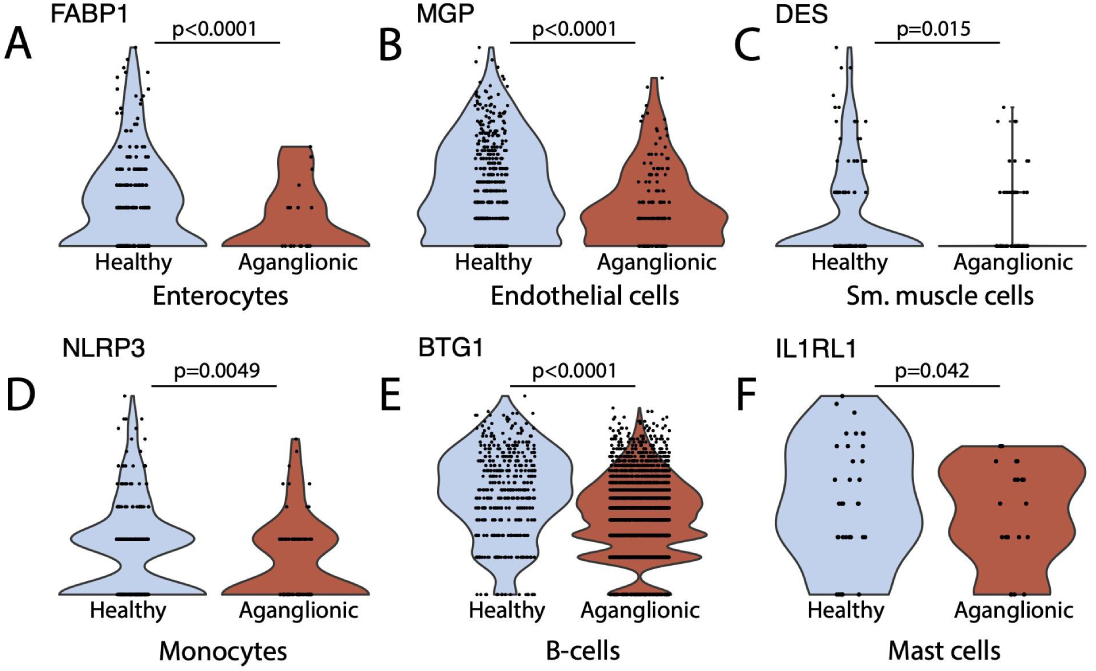
Expression of key DEG genes in healthy and aganglionic colon specimens. (Wilcoxon rank sum test, min.diff.pct = 0.25, logfc.threshold = 0.25).

**Table 1:** List of downregulated genes in aganglionic colon residing cell populations of HSCR patients based on scRNA-seq analysis. Genes that were found significantly downregulated in all three HSCR patients in corresponding cell types (Wilcoxon rank sum test, min.diff.pct = 0.25, logfc.threshold = 0.25).

| Cell type | Downregulated genes in aganglionic colon of HSCR patients |
| --- | --- |
| Enterocytes | *FABP1* |
| Goblet cells | *FABP1* |
| Smooth muscle cells | *DES* |
| Endothelial cells | *MGP*, *DEPP1*, *F2RL3* |
| Fibroblasts | *CFD*, *APOE* |
| Monocytes | *AREG*, *HSP9081*, *MT2A*, *GPR183*, *EZR*, *NLRP3*, *ITM2C* |
| Mast cells | *FTH1*, *EIF1*, *GLUL*, *NR4A2*, *H3F3B*, *ZFAND5*, *MAP1LC3B*, *IER2*, *CDKN1A*, *POLR2L*, *SOCS1*, *ID2*, *TRA2A*, *NEAT1*, *ANKRD28*, *CTSG*, *VMP1*, *IL1RL1*, *AC020571.1* |
| B-cells | *BTG1*, *SLC2A3* |
| T-cells | *ZFP36L2*, *GPR183*, *LTB*, *RSRP1*, *JUNB*, *EEF1A1*, *CCNL1*, *SLC2A3* |

We hypothesize that the altered transcriptomic profile of the aganglionic colon tissue can lead to hindered neural progenitor cell differentiation. Notably, mast cells - which are key regulators of epithelial permeability, neuroimmune interactions, and peristaltic movements (29) - showed significant downregulation of a number of genes, such as *IL1RL1*, which is involved in their activation (**Fig. 3F**). Activated mast cells produce growth factors and other mediators contributing to neuroimmune communication and influencing neural cell differentiation (30). Downregulation of IL1RL1 may suggest that the lack of mast cell activation could play a role in the impaired differentiation of neural progenitor cells in the aganglionic colon.

Additionally, we identified significant alterations in the expression of other key genes that may be important in HSCR, potentially contributing to the disease’s characteristic symptoms, such as disrupted gut motility, altered epithelial function, and immune dysregulation. We observed a significantly lower expression of Liver Fatty Acid Binding Protein 1 (FABP1) in cells of epithelial origin (enterocytes, goblet cells) (**Table 1 and Fig. 3A**). *FABP1* is a key marker gene of colonocytes, a specific intestinal epithelial cell type (26) that has an important function in regulating the gut microbiome (31). We have identified significant downregulation of several genes in aganglionic-segment endothelial cells including *MGP*, a gene coding the GLA matrix protein (MGP) (**Fig. 3B**), that has been shown to modulate the activity of bone morphogenetic protein-2 (BMP-2) (32, 33). In aganglionic-colon resident smooth muscle cells, significantly lower *DES* desmin gene expression levels were observed compared to the healthy colon (**Table 1 and Fig. 3C**) in good correlation with earlier immunofluorescence stainings of human colon samples that showed decreased expression of desmin in the smooth muscle cells of the aganglionic colon segment of HSCR patients (34).

A substantial number of genes were downregulated in immune cells residing in the aganglionic colon, including T-cells, B-cells and monocytes (**Table 1**). Notably, reduced expression of *BTG1* was observed in B-cells (**Fig. 3E**), and *NLRP3* was decreased in monocytes (**Fig. 3D**). These transcriptional alterations are similar to those reported in inflammatory bowel diseases (35, 36), suggesting a potential role for immune cell dysregulation in the pathogenesis of HSCR.

### Germline mutations predominate in HSCR pointing to a limited role for somatic variants

Based on our findings, hindered differentiation of neural progenitor cells might be responsible for the lack of mature neurons in the aganglionic colon due to transcriptomic changes in the progenitor cells and surrounding stroma cells (**Fig. 2 and 3**). Using WGS data we aimed to identify associated genomic alterations in the aganglionic colon that can lead to deficient neural progenitor cells differentiation.

As previously noted, we identified several germline variants in key HSCR-associated genes; however, no aganglionic colon-specific alterations were detected in known HSCR-associated genes. Moreover, in receptor genes of several signaling pathways (e.g. *NOTCH* genes, *SMO* and *PTCH1, BMRP1*) germline mutations have been identified in several patients (**Supplementary table 3**). We have observed missense germline variants in *BMP4 (*patient 1 and 3) and *JAG1* (patient 2), genes regulating the BMP and NOTCH signaling pathways, however, we did not find statistically significant differences between their expression in healthy and aganglionic colon stroma cells (data not shown).

On the other hand, the number of somatic variants were generally low in our patient cohort (**Fig. 4A**) with a small number of aganglionic colon-specific variants identified. Analysis of the allele frequencies revealed that most of these variants were subclonal heterozygous mutations (**Fig. 4B**). Despite the limited number of aganglionic-specific variants, we detected somatic mutations in transcription factor binding sites of *TEAD4, RFX5* and *HOXB* (**Fig. 4C**). It is important to note that sequencing at 60x depth in HSCR patient samples poses significant challenges for detecting somatic variants present in low-frequency cell populations, limiting our ability to pinpoint critical somatic alterations potentially driving impaired neural differentiation in HSCR.

**Figure 4:**
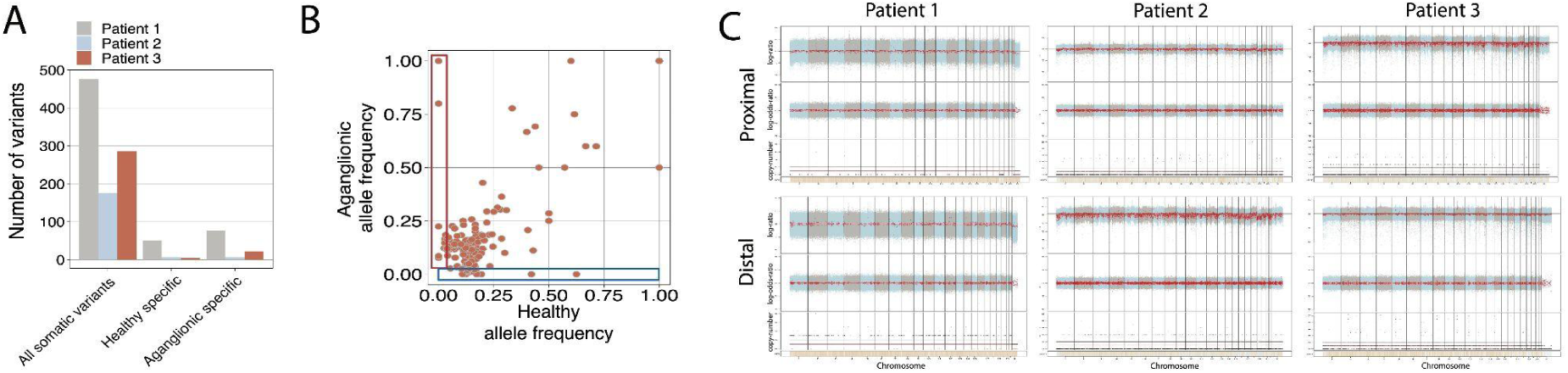
Genomic analysis of HSCR patient samples. (**A**) Number of identified somatic variants and healthy- or aganglionic colon segments specific variants in three HSCR patients. (**B**) Representative allele frequency plot of identified colon specific variants showing healthy colon specific (blue) and aganglionic colon specific (red) variants. (**C**) CNV analysis on three HSCR patients using FACETS. Plots show copy number state (bottom), log-odds ratio (middle) and log-ratio (top) for healthy- (top row) and aganglionic (bottom row) samples.

We performed copy number variation (CNV) analysis in healthy and aganglionic colon samples. FACETS analysis did not reveal any chromosomal-level CNV events in the aganglionic regions across all three patients studied (**Fig. 4D**).

## Discussion

In this study, we employed single-cell RNA sequencing (scRNA-seq) and whole genome sequencing (WGS) on human colon samples from HSCR patients to comprehensively characterize transcriptomic and genomic alterations associated with the disease.

scRNA-seq enabled the identification of 13 distinct cell types, including minor populations of neurons, glial cells, and neural progenitor cells (**Fig. 1E**). Consistent with the known pathology of HSCR, mature neurons were absent in the aganglionic colon segments; however, we unexpectedly observed the presence of neural progenitor cells in both healthy and aganglionic regions across all patient samples (**Fig. 1F**, **Fig. 2B**). These progenitor cells expressed high levels of specific marker genes such as *NGFR* and *PTN* (**Fig.2C**). Compared to mature neurons, neural progenitor cells displayed reduced expression of several HSCR-associated genes, including *ECE1, ZEB2, GFRA1, SEMA3D* and *TCF4* (**Fig. 2C**). Additionally, neural progenitor cells in the aganglionic colon exhibited downregulated expression of receptor genes involved in key signaling pathways, such as *NOTCH2, BMPR1A, FGFR1, RXRA* and *RXRB*, relative to progenitor cells in the healthy colon (**Fig. 4A**). There is only a very limited number of studies that postulated the presence of neural progenitor cells in HSCR patients’ distal colon. Wilkinson et al. have identified neural progenitor cells in the aganglionic colon of HSCR patients (37). They showed that these progenitor cells can differentiate to mature neurons *in vitro* and show normal migratory behavior in mouse gut explants (37). These data together support the hypothesis that neural progenitor cell differentiation is specifically hindered in the aganglionic colon, most probably due to transcriptomic changes in the progenitor cells and surrounding stroma cells.

Through collected scRNA-seq data of three HSCR patients we have been able to identify human colon resident cell types (**Fig. 1**) and perform differential gene expression analysis for each cell type to better understand the transcriptomic changes observed in HSCR (**Table 1**). In colon epithelial cell-types including enterocytes and goblet cells decreased *FABP1* (fatty acid-binding protein 1) expression has been observed (**Table 1** and **Fig. 3A**). *FABP1* is essential for proper lipid metabolism in differentiated enterocytes (38), particularly concerning fatty acid uptake and it is also a marker gene of early enterocytes (39). *FABP1* is involved in the regulation of gene expression, cell growth and proliferation and modulation of signaling pathways (40). *FABP1* has a role in the regulation of inflammation and has been linked with the immune microenvironment of certain colorectal carcinomas (40, 41). In human colon organoids, FABP1 expression has been increased in the presence of FGF and IGF-1 differentiation factors (39) however in HSCR patients we did not find significant differences between FGF or IGF-1 expression of the healthy and aganglionic colon segments (data not shown). This observation in combination with other transcriptomic changes in aganglionic colon resident endothelial cells (*MGP*, **Fig. 3B**) and smooth muscle cells (*DES*, **Fig. 3C**) along with transcriptional changes in the immune cell repertoire (**Fig. 3D-F**) shows that HSCR leads to a wide range of transcriptomic changes not only in the neural subsets, but other cell types as well. Some of these expression changes might be important in embryonic development and progenitor differentiation.

Recently, connection between the pathogenesis of HSCR and immune cells, especially mast cells has been postulated by various studies. In the aganglionic colon segment of HSCR patients, an increased number of mast cells was identified by several studies (42, 43), however in our patient samples we did not observe elevated mast cell numbers in the aganglionic colon (**Fig. 1F**). Mast cells form close contacts with nerve fibers in the aganglionic colon segment of HSCR patients and, by producing NGF, they might have a role in the regulation of differentiation and proliferation of neural cells (42, 44). Using our scRNA-seq data from HSCR patients we identified mast cell populations in our patients and compared the gene expression profile of healthy- and aganglionic colon resident mast cells and identified a large number of genes that were downregulated in the aganglionic colon resident mast cells (**Table 1**). Among the significantly downregulated genes we could identify *SOCS1* (suppressor of T-cell signaling 1), a negative regulator of c-Kit and interleukin-3 (IL-3) receptor signaling (45), *IL1RL1*, that encodes the receptor for IL-33 (46), *FTH1*, the gene coding ferritin, the main iron-binding intracellular protein. Ferritin is known to have direct modulatory activity on T cells and mast cell released ferritin particles may help to moderate inflammation at local sites (47). Moreover, ferritin has been described to have major roles in the regulation of iron homeostasis and has been linked with neurodegenerative diseases (48). Taken together, based on collected scRNA-seq we can conclude that mast cells in HSCR show significant transcriptomic dysregulation of multiple genes that might influence mast cell activity and their interactions with neurons and neural progenitor cells.

HSCR is a genetic neurodevelopmental disease, and more than 20 genes have been identified that play a role in its development (49). In fact, we have demonstrated the presence of several missense or synonymous mutations in well-known HSCR-associated genes including *RET, DHCR7, EDNRB, SEMA3D* or *NRG1/3* in our cohort (**Supplementary table 2**). Furthermore, we have identified several germ-line variants in NOTCH, BMP, FGF and Hedgehog signaling pathway genes as well (**Supplementary table 3**). However, such germ-line variants alone can not explain transcriptomic differences found between the healthy and aganglionic colon segment specific cells (**Table 1** and **Fig. 2E**). The complex inheritance of HSCR and the fact that only 30% of HSCR patients harbor known pathogenic germ-line variants (17) raises the possibility of the involvement of somatic variants in HSCR pathogenesis. However, most HSCR studies aiming to identify somatic colon-specific variants focused on RET gene variants (50)(51)(52) and no pathogenic somatic variants were identified in analyzed patients. By performing WGS on the healthy and aganglionic colon segments in our cohort we were able to identify a small number of colon-specific and aganglionic colon segment specific variants (**Fig. 4B, C**). We showed that the number of somatic variants is low in HSCR patient samples and only a very limited number of aganglionic colon specific variants were identified including somatic variants in transcription-factor binding sites of TEAD4, RFX5 and HOXB (**Fig. 4C**) and various intronic and upstream gene variants. Previously, TEAD4 activity has been shown to be significantly altered in short segment HSCR cases (53). By performing CNV calling on the patient cohort we did not find significant CNV events either in healthy or aganglionic colon segments (**Fig. 4D**). Together these results suggest that similarly to the findings of other research groups, somatic variations or local CNV events do not play a major role in the pathogenesis of HSCR, but the combination of pathogenic germ-line variants with non-coding- and somatic variants along with transcriptomic changes together lead to symptomatic disease. However, we understand that the applied sequencing strategy is not suitable to find low frequency, cell-type or tissue specific somatic variants that might play crucial roles in HSCR. For the identification of such variants pure, isolated cell populations and deep sequencing might be required.

To our knowledge, we are the first to use scRNA-seq on HSCR patient samples to compare the transcriptome of the healthy and aganglionic colon at a single-cell level. scRNA-seq analysis of HSCR colon samples demonstrated that neural progenitor cells are present in the aganglionic colon segment of HSCR patients, which raises the possibility that the aganglionic colon can be possibly used for future autologous transplantation therapies and to design therapeutic modalities based on the stimulation of neurogenesis in the aganglionic colon region *in vivo*. Our transcriptomic analysis highlighted recurrent gene expression changes in HSCR patients that might be important in the pathogenesis of HSCR. Combined with the performed genomic analysis our results underline the importance of gene expression changes in the etiopathogenesis of HSCR.

## Methods

### Tissue samples

We queried for patients diagnosed with Hirschsprung’s disease between 25th July 2019 and 28th October 2020 through the Hirschsprung Disease Research Collaborative. All enrolled patients had a short segment (aganglionosis of the rectum or rectosigmoid colon) Hirschsprung’s disease without trisomy 21 as it was validated by a board-certified pathologist. All individuals were ascertained with written informed consent approved by the Institutional Review Board of the enrolling institution. From each patient, healthy (proximal), aganglionic (distal) colon tissue samples and blood samples were collected. In our study was not considered as a biological variable.

### Tissue dissociation

Surgically resected tissue was collected in RPMI 1640 medium (Gibco). The sample is minced into small pieces (∼2 mm). The pieces were put into a Miltenyi C-tube with 4.75 ml of RPMI 1640 medium with 250 *μ*l of 100 mg/ml dispase/collagenase mixture (Roche 11-097-1130-001). The C-tube is placed on the gentleMACS Dissociator and the program m_intestine_01 was executed. The C-tube was attached to the MACSmix Tube Rotator for 30 min at 37°C. This procedure was repeated again, and followed by one final agitation cycle performed on the gentleMACS Dissociator. The cell suspension was applied to a 70 *μ*m MACS SmartStrainer and washed with 5 ml of RPMI 1640 into a 15 ml conical tube. The cells were counted to determine viability using 0.4% Trypan blue on a Countess II (Thermo Fisher). Cells were centrifuged at 300×g for 5 min at 4°C. The cells were resuspended in cryopreservation media (50% RPMI 1640, 40% FBS, 10% DMSO) and cryopreserved prior to sequencing. Cell viability was assessed prior to 10X Genomics preparation for droplet sequencing.

### Immunostaining

Paraffin sections were dewaxed in Hemo-De, rehydrated in a decreasing ethanol series and washed in water and PBS. Endogenous peroxidase activity was blocked with 3% H_2_O_2_ and antigen retrieval was performed by heating sections in 10 mM sodium citrate buffer (pH 6.0) at 95°C for 10 min in a pressure cooker. After washes in PBS and water, the sections were blocked in 10% donkey serum in 10mM sodium citrate buffer for 1 hr and then probed with primary antibody (p75, 1:200, Abcam, ab52987) overnight. For immunofluorescence staining, the sections were incubated with corresponding fluorescent labeled secondary antibody Alexa Fluor 647 donkey anti-Rabbit IgG (1:500, Thermo Fisher, A-31573) for 1 hr, counterstained with DAPI (Invitrogen, R37606) and mounted with ProLong diamond antifade mountant (Invitrogen, P36970). Images were taken on Zeiss 700 confocal inverted microscope with x20 objective and merged in FIJI (NIH ImageJ).

### Single cell library preparation and sequencing

Following tissue dissociation of healthy and aganglionic colon samples, live cells were washed twice with PBS + 0.04% BSA to remove ambient RNA. Cells were loaded onto the Chromium Next GEM Chip (10x Genomics) for each donor. GEM generation and barcoding, reverse transcription, cDNA generation and library construction were performed following the manufacturer’s protocol (Single cell 3′ reagent kit v.3.0, 10x Genomics). Single-cell libraries were pooled and sequenced in paired-end reads on Novaseq (Illumina).

### Single cell RNA sequencing data analysis

Clustering, differential gene expression analysis was performed using the *Seurat* package (54). Low abundant genes (expressed in less than 3 cells) and cells of potentially low quality (total UMI<700 or total UMI>15,000, or percentage of mitochondrial genes >25%) were removed from downstream analysis. For normalization of our datasets we used *SCtransform* (55) and for dimensional reduction and clustering we used *Seurat* (54). 3,000 highly variable genes were selected in each sample. Anchors between individual data were identified and correction vectors were calculated to generate an integrated expression matrix, which was used for subsequent clustering. Integrated expression matrices were scaled and centered followed by principal component analysis (PCA) for dimensional reduction. PC1 to PC20 were used to construct nearest neighbor graphs in the PCA space followed by Louvain clustering to identify clusters (resolution=0.5). For visualization of clusters, Uniform Manifold Approximation and Projection (UMAP) was generated using PC1 to PC20. To identify markers (differentially expressed genes) for clusters and assigning cell type identity of clusters, expression value of each gene in given clusters were compared against the rest of cells using Wilcoxon rank sum test and p-values were adjusted with the number of genes tested. Gene with log fold change >0.25 and adjusted p-value <0.05 were considered to be significantly enriched. Using canonical markers of human colon-specific cell types, cell identities were assigned to clusters.

### Whole genome sequencing and data analysis

WGS data with 60X coverage was collected from three HSCR patients’ healthy- and aganglionic colon and corresponding blood samples. Genomic DNA was extracted from fresh frozen tissue sections along with whole blood using Qiagen’s DNeasy Blood and Tissue kit (Qiagen). Libraries were prepared using the Nextera DNA Flex Library Prep Kit (Illumina). Libraries were sequenced on a NovaSeq 6000 instrument targeting 300 million read-pairs on a 2 x 150 bp run.

FASTA files were generated from paired-end sequencing and aligned to the HG38 reference genome using *BWA-MEM* (56). Default settings were used during alignment. *Samblaster* was used to remove duplicate reads from the output generated by the alignment (57). *Samtools merge* was used to merge BAM files when a sample was run in multiple lanes. Finally, *Samtools sort* was used to sort all BAM files by leftmost coordinates (58).

*Freebayes* was used to call variants for each patient (59). For variant annotation we used *Bedtools intersect* and removed any called variants that did not fall within exonic regions (60). We also removed variants within low complexity regions to avoid bias due to the highly repetitive nature of these regions. Using *Snpsift filter*, we filtered out variants with a quality score less than or equal to 20 (61). Variants with a sample depth of 50 or less were also removed. *VT view* was used to identify both biallelic and multiallelic variants (62). Only biallelic variants were used in our analysis. To annotate and predict the effects of each variant, we used *Snpeff* (62, 63). *Snpsift filter* was then used to remove any variants that had more than 5 alternate allele observations (AO) in the control sample (collected blood), to remove germline variants and leave only somatic variants. Using the same filtering criteria on the somatic variants we identified variants specific to aganglionic-colon samples as well. Copy number variants were called using the *FACETS* R package (64). Larger variants were called with RUFUS.

### Statistical analysis

For the comparison of two samples from normally distributed populations with equal variances, Student’s *t*-test was performed, while in case of unequal variances a Welch’s t-test was applied. The assumptions of normality and equal variance were checked by Kolmogorov-Smirnov test. Multiple comparisons were performed with analysis of variance (ANOVA). A p-value cutoff of 0.05 was used as the threshold of significance.

### Study approval

All patients provided informed consent for the collection of human specimens and it was approved by the University of Utah Institutional Board in accordance with the U.S. Common Rule.

## Data availability

The processed datasets generated and/or analyzed in the study and the supplementary information are available in the GitLab repository of the manuscript: https://gitlab.com/tariszabi/hscr_study

## Acknowledgement

S.T., A.F., Y.Q., X.H., and G.T.M. were supported by NIH grants U24 CA209999 and R01 HG009000. G.T.M. was supported by NCI award K23 CA212271. Research reported in this publication utilized the High-Throughput Genomics and Bioinformatic Analysis Shared Resource at Huntsman Cancer Institute at the University of Utah and was supported by the National Cancer Institute of the National Institutes of Health under Award Number P30 CA042014. The support and resources from the Center for High-Performance Computing at the University of Utah are gratefully acknowledged. The NIH Shared Instrumentation Grant 1S10OD021644-01A1 partially funded the computational resources used for this study. We are grateful to the patients who provided tissue samples for these studies to the Colorectal Center at Primary Children’s Hospital. The content is solely the responsibility of the authors and does not necessarily represent the official views of the National Institutes of Health.

## Authors contributions

S.T. contributed to the genomic data processing and analysis, variant identification, transcriptomic analysis, results interpretation, visualization. X.H. and Y.Q. contributed to the study design, project coordination, interpretation. M.R. obtained patient tissue. A.F. contributed to genomic data processing and analysis. A.L. performed histological stainings and validation of patient status. L.M., A.L, T.M. and P.J.M. contributed to patient sample selection, DNA extraction and tissue dissociation. G.K. and M.A.F. performed the immunofluorescence stainings and fluorescent microscopic experiments. X.H., Y.Q., M.R. and G.T.M conceived the project. S.T., X.H., Y.Q., M.R. and G.T.M. contributed to manuscript writing. All authors contributed to the manuscript refinement. All authors read and approved the final manuscript.

