## Supplementary_data for "Single-cell RNA sequencing in Hirschsprung’s disease tissues reveals lack of neuronal differentiation in the aganglionic colon segment"

Institute of Human Genetics

University of Utah School of Medicine

15 North 2030 East, Room 7410B Salt Lake City, UT 84112-5330

The authors have declared that no conflict of interest exists.

### SUPPLEMENTARY DATA

**<u>Supplementary Table 1:</u> List of identified cell types in scRNAseq analysis and associated marker genes**

| **Cell type** | **Marker Genes:** |
|---|---|
| Enterocytes | *EPCAM+, KRT8+, KRT18+* |
| Goblet cells | *TFF3+, MUC2+, FCGPB+* |
| Endothelial cells | *PECAM1+, VWF+, CD93+* |
| Tuft cells | *TRPM5+, DCLK1+, GFI1B+* |
| Fibroblasts | *COL1A2+, COL3A1+, CFD+* |
| Smooth muscle cells | *ACTA2+, MYH11+, RGS5+* |
| B-cells | *IGHM+, CD79A+, CD83+, CXCR4+* |
| T-cells | *CD3D+, IL7R+, CD7+, CD2+* |
| Monocytes | *CD14+, LYZ+, IL1B+* |
| Mast cells | *KIT+, TPSAB1+, TBSB2+* |
| Neural progenitor cells | *NGFR+, UCHL1-, S100B-* |
| Mature neurons | *TUBB3+, UCHL1+, GAL+* |
| Mature glial cells | *S100B+, ERBB3+* |

**<u>Supplementary Table 2:</u> List of germline variants in HSCR-associated genes of HSCR patients**

Accession numbers, status and condition is based on the ClinVar database. For variants that are not present in ClinVar, nucleotide change is provided (missense mutations).

| | Gene | Type | Accession # | Status | Condition |
|---|---|---|---|---|---|
| **Patient 1** | DHCR7 | Missense | VCV000093708 | Benign | - |
| | RET | Synonymous | VCV000095995 | Benign | - |
| | | Synonymous | VCV000167590 | Benign | - |
| | SEMA3D | Missense | - | - | c.2101A>C |
| | NRG1 | Missense | - | - | c.866T>C |
| | NRG3 | Synonymous | VCV000691408 | Uncertain | Aganglionic megacolon |
| | EDNRB | Synonymous | VCV000226622 | Benign | - |
| | | Synonymous | VCV000226625 | Benign | - |
| **Patient 2** | SEMA3D | Missense | - | - | c.2101A>C |
| | RET | Missense | VCV000024934 | Conflicting pathogenicity | Hirschsprung's disease, Multiple endocrine neoplasia |
| | KIF1BP | Missense | - | - | c.196G>A |
| | DHCR7 | Missense | VCV000093707 | Benign | Smith-Lemli-Opitz syndrome, neurodevelopmental disorders |
| | | Missense | VCV001177469 | Benign | Smith-Lemli-Opitz syndrome, neurodevelopmental disorders |
| | TCF4 | Missense | VCV000160083 | Benign | Pitt-Hopkins Syndrome, Corneal dystrophy |
| **Patient 3** | DHCR7 | Splice acceptor | VCV000093725 | Pathogenic | Smith-Lemli-Opitz syndrome, neurodevelopmental disorders |
| | | Missense | - | - | c.659T>C |
| | | Missense | VCV001177469 | Benign | Smith-Lemli-Opitz syndrome |
| | SEMA3D | Missense | - | - | c.2101A>C |
| | KIF1BP | Missense | - | - | c.196G>A |
| | NRG3 | Missense | VCV000691406 | Uncertain | Aganglionic megacolon |
| | GFRA1 | Missense | VCV001328035 | Benign | - |

**<u>Supplementary table 3:</u> List of germline variants in various signaling pathway receptor genes of HSCR patients**

Accession numbers, status and condition is based on the ClinVar database.

| | Gene | Type | Accession # | Status | Condition |
|---|---|---|---|---|---|
| **Patient 1** | NOTCH2 | Missense | VCV000134972 | Benign | Hajdu-Cheney Syndorme |
| | | Missense | VCV000256152 | Benign | Cerebral arteriopathy, Lateral meningocele syndrome |
| | | Missense | VCV001181077 | Benign | Cerebral arteriopathy, Lateral meningocele syndrome |
| | | Missense | VCV000811011 | Benign | Cerebral arteriopathy, Lateral meningocele syndrome |
| | NOTCH4 | Missense | VCV001297197 | Benign | - |
| | | Missense | VCV001287249 | Benign | - |
| | | Missense | VCV001294880 | Benign | - |
| | PTCH1 | Missense | VCV000041663 | Benign | Hereditary cancer-predisposing syndrome, Gorlin syndrome |
| | BMPR1A | Missense | VCV000041782 | Benign | Juvenile polyposis syndrome |
| **Patient 2** | NOTCH4 | Missense | VCV001220553 | Benign | - |
| | | Missense | VCV001297197 | Benign | - |
| | | Missense | VCV001287249 | Benign | - |
| | | Missense | VCV001294880 | Benign | - |
| | NOTCH3 | Missense | VCV000256152 | Benign | Cerebral arteriopathy, Lateral meningocele syndrome |
| | | Missense | VCV001181077 | Benign | Cerebral arteriopathy, Lateral meningocele syndrome |
| | | Missense | VCV000811011 | Benign | Cerebral arteriopathy, Lateral meningocele syndrome |
| **Patient 3** | PTCH1 | Missense | VCV000041663 | Benign | Hereditary cancer-predisposing syndrome, Gorlin syndrome |
| | BMPR1A | Missense | VCV000041782 | Benign | Juvenile polyposis syndrome |
| | NOTCH3 | Missense | VCV000256152 | Benign | Lateral meningocele syndrome, cerebral arteriopathy |
| | | Missense | VCV001181077 | Benign | Lateral meningocele syndrome, cerebral arteriopathy |
| | | Missense | VCV000811011 | Benign | Lateral meningocele syndrome, cerebral arteriopathy |

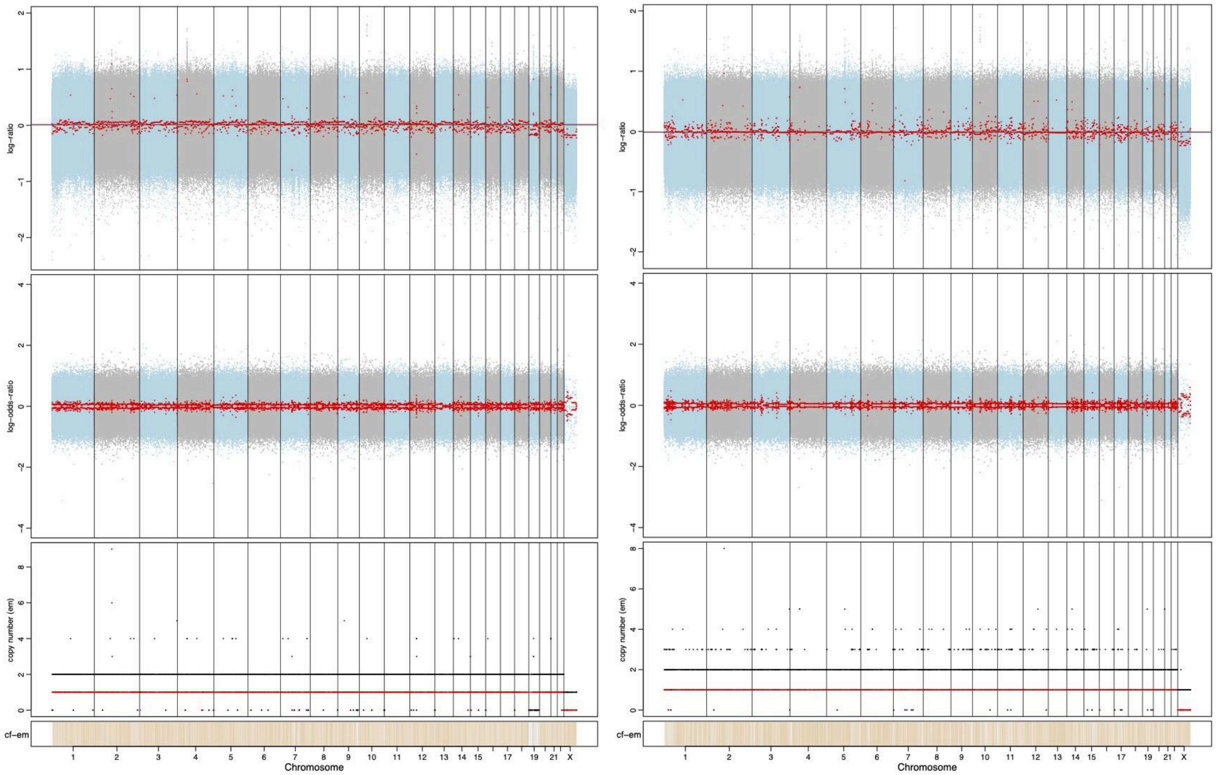


**Supplementary Figure 1:** FACETS output of Patient 1 healthy (left) and aganglionic (right) colon samples.

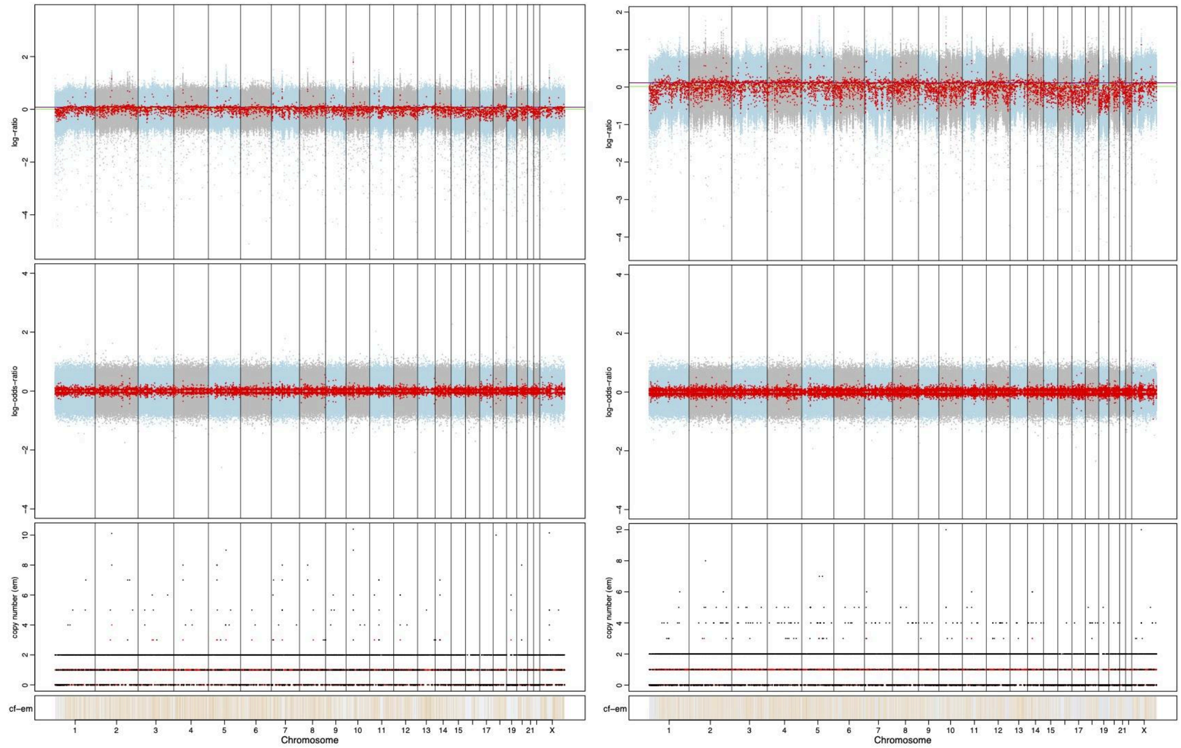


**Supplementary Figure 2:** FACETS output of Patient 2 healthy (left) and aganglionic (right) colon samples.

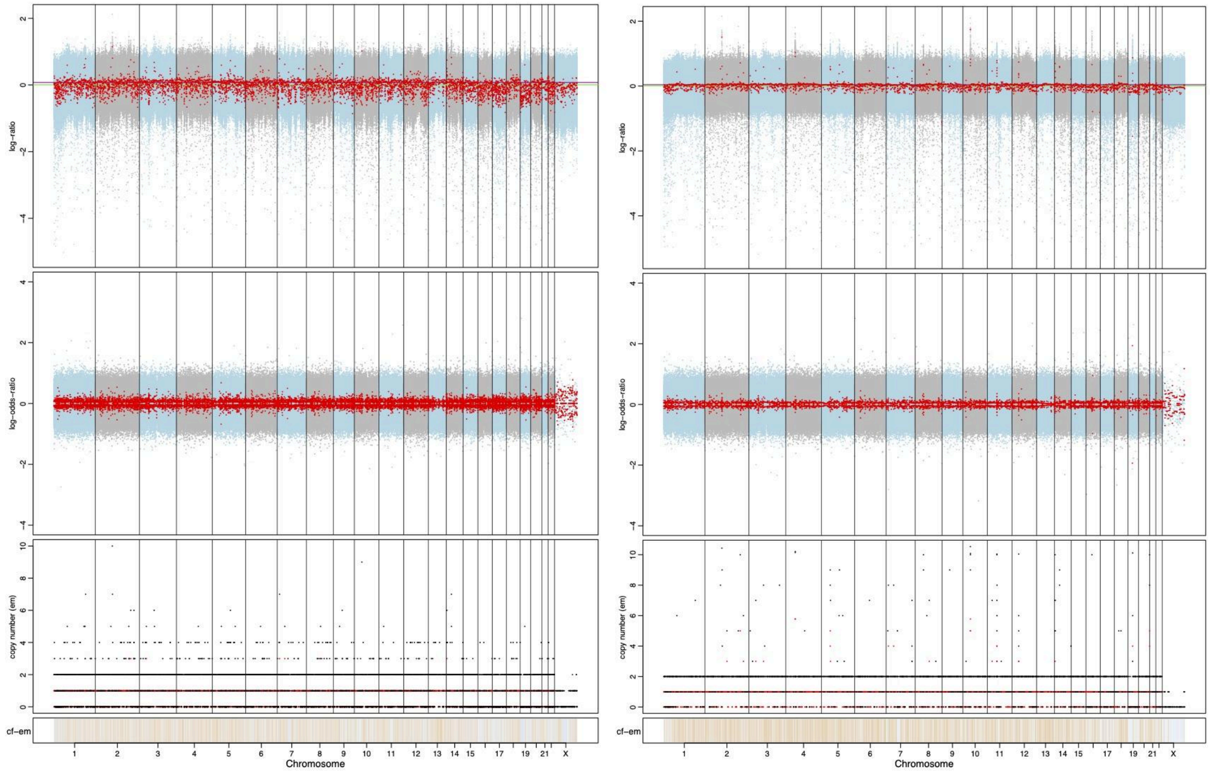


**Supplementary Figure 3:** FACETS output of Patient 3 healthy (left) and aganglionic (right) colon samples.

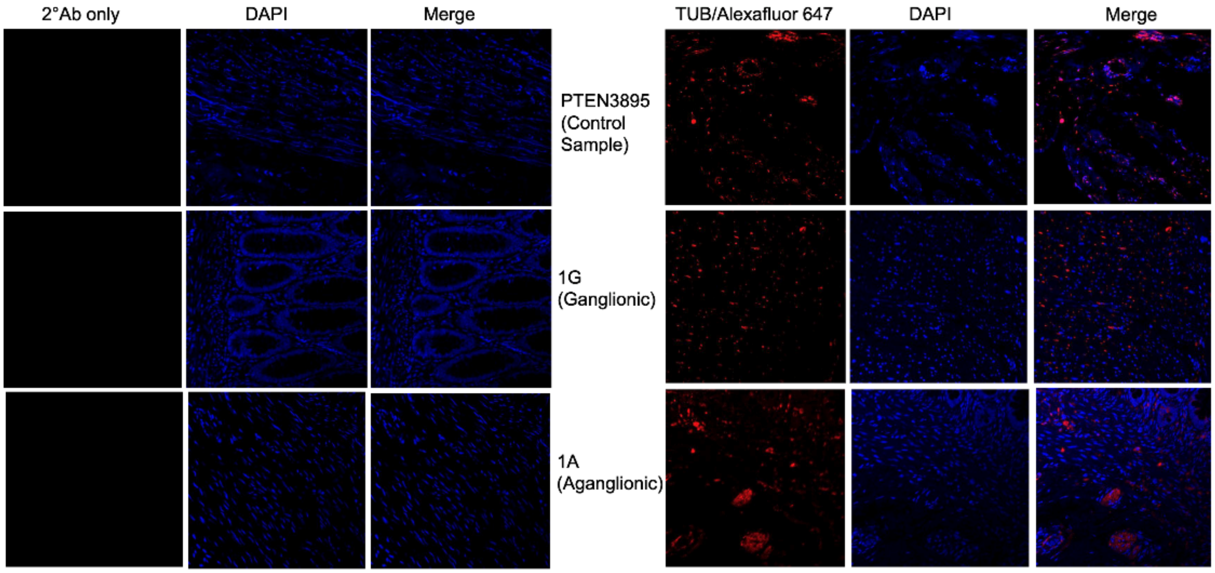


**Supplementary Figure 4:** Representative images from fluorescence microscopic analysis of HSCR healthy and aganglionic colon segments using DAPI and anti-p75 (*NGFR*) antibody staining. As positive control, PTEN3895 cells were used.
